# H3K27 Acetylated Nucleosomes Facilitate HMGN Protein Localization at Regulatory Sites to Modulate The Interaction of Transcription Factors with Chromatin

**DOI:** 10.1101/2021.05.03.442466

**Authors:** Shaofei Zhang, Yuri Postnikov, Alexei Lobanov, Takashi Furusawa, Tao Deng, Michael Bustin

## Abstract

Nucleosomes containing acetylated H3K27 are a major epigenetic mark of active chromatin and identify cell-type specific chromatin regulatory regions which serve as binding sites for transcription factors. Here we show that the ubiquitous nucleosome binding proteins HMGN1 and HMGN2 bind preferentially to H3K27ac nucleosomes at cell-type specific chromatin regulatory regions. HMGNs bind directly to the acetylated nucleosome; the H3K27ac residue and linker DNA facilitate the preferential binding of HMGNs to the modified nucleosomes. Loss of HMGNs increases the levels of H3K27me3 and the histone H1 occupancy at enhancers and promoters and alters the interaction of transcription factors with chromatin. These experiments indicate that the H3K27ac epigenetic mark affects the interaction of architectural protein with chromatin regulatory sites and provide insights into the molecular mechanism whereby ubiquitous chromatin binding proteins, which bind to chromatin without DNA sequence specificity, localize to regulatory chromatin and modulate cell-type specific gene expression.

## INTRODUCTION

Correct binding of transcription factors (TFs) to their specific DNA motifs in chromatin plays a key role in establishing an epigenetic landscape that facilitates cell-type specific gene expression necessary for the maintenance of cell identity (D’Alessio et al. 2015; Lambert et al. 2018). Dysregulation of the interaction of TFs with chromatin can lead to changes in gene expression and destabilize cell identity, thereby leading to disease. The interaction of TFs with chromatin is dynamic; TFs continuously move throughout the nucleus and reside temporarily at their specific binding sites (Phair et al. 2004; Voss and Hager 2014; Zaret et al. 2016). Binding of TFs to chromatin is facilitated by nuclear factors that help TFs access their binding sites and perturbed by factors that impede the binding of TFs to specific chromatin sites, especially at enhancers and promoters (Klemm et al. 2019), chromatin regions enriched in H3K27ac nucleosomes (Hnisz et al. 2013). Possible regulators of TF chromatin binding include architectural proteins such as the linker histone H1 and high mobility group (HMG) proteins, these ubiquitous structural proteins that are known to affect chromatin compaction and organization in many cell-types. Histone H1 facilitates the formation of higher order chromatin organization and stabilizes compact chromatin structures that inhibit transcription (Hergeth and Schneider 2015; Jordan 2016; Fyodorov et al. 2018), while HMG proteins are mostly associated with reduced chromatin compaction and increased gene expression from chromatin templates (Bustin 2010; Reeves 2010). Given the global effects of H1 and HMGs on chromatin structure and gene expression, it is likely that these ubiquitous structural proteins do modulate the binding of TFs to chromatin, a possibility that has not been studied in detail. Here we focus on the high mobility group N (HMGN) protein family and show that the major members of this family, HMGN1 and HMGN2, bind preferentially to nucleosomes containing the H3K27ac epigenetic mark, and affect the binding of TFs to chromatin.

HMGN is a family of structural proteins that bind to nucleosomes without specificity for the underlying DNA sequence (Postnikov and Bustin 2010). The amino acid sequence of HMGN1 and HMGN2 is evolutionarily conserved and these proteins have been detected in every vertebrate cell examined (Gonzalez-Romero et al. 2015). Live cell imaging experiments revealed that the interaction of HMGN proteins with chromatin is highly dynamic; HMGNs bind to nucleosomes with short residence times and can be readily dislocated from their chromatin binding sites (Catez and Hock 2010). The binding of HMGNs to nucleosomes reduces chromatin compaction, most likely because it alters the interactions of linker histone H1 with chromatin (Catez et al. 2002; Rochman et al. 2009; Murphy et al. 2017) and because it could reduce access to the nucleosome acidic patch, a region thought to facilitate and stabilize interactions between neighboring nucleosomes (Kato et al. 2011; Kalashnikova et al. 2013). Although HMGNs bind to chromatin without DNA sequence specificity, they preferentially localize to chromatin regulatory sites that are easily digested with DNase I and are enriched in H3K27ac modified histones, such as enhancers and promoters (Deng et al. 2015; He et al. 2018). Numerous studies show that changes in HMGN levels are frequently associated with a wide range of cell-type specific changes in gene expression and phenotypes (Nanduri et al. 2020). Genetically altered mice lacking both HMGN1 and HMGN2 proteins (DKO mice) are born and survive but show multiple phenotypes (Deng et al. 2015), (https://www.mouseclinic.de/). MEFs isolated from DKO mice can be reprogramed into pluripotent cells by exogeneous TFs more efficiently than MEFs isolated from WT mice suggesting that loss of HMGNs destabilized the maintenance of cell identity (He et al. 2018; Garza-Manero et al. 2019). In Down syndrome, one of the most prevalent human genetic diseases, the presence of an extra copy of *HMGN1* has been directly linked to increased levels of H3K27ac and to gene expression changes that may lead to increased incidence of acute lymphoblastic leukemia (Lane et al. 2014; Cabal-Hierro et al. 2020). Taken together, the available data suggest that HMGNs modulate and fine tune cell-type specific gene expression programs.

Given that the amount of HMGN in a cell is enough to bind only about 1% of the nucleosomes (Bustin 1999), it is not clear how these structural proteins, that interact with chromatin without DNA-sequence specificity, can nevertheless affect gene expression and cellular phenotypes in a cell-type specific manner. Most likely, HMGNs affect cell-type specific gene expression by preferentially localizing to chromatin regulatory regions, as indicated by genome-wide analyses that show high HMGN occupancy at chromatin regions containing high levels of H3K27ac (Deng et al. 2015; He et al. 2018), an epigenetic modification that marks transcription start sites and enhancers (Strahl and Allis 2000; Heintzman et al. 2009; Hnisz et al. 2013). Conceivably, the presence of HMGN at enhancers and promoters may affect the interaction of TFs with these sites, thereby impacting cell-type specific gene expression. The mechanism that targets HMGN to enhancers and promoters and the possible effect of these proteins on TFs chromatin binding have not yet been investigated.

Here we delineate the mechanism that facilitates the preferential binding of HMGN proteins to chromatin regulatory sites and examine whether HMGNs affect the binding of transcription factors to chromatin. Using bioinformatic analyses we demonstrate that HMGN1 and HMGN2 preferentially localize to nucleosomes containing the H3K27ac residue. We show that the presence of H3K27ac, an epigenetic mark of active chromatin, strengthens the binding of both HMGN1 and HMGN2 to the modified nucleosomes, and that loss of HMGNs alters H3K27 modifications and H1 occupancy at enhancers and promoters. We analyze the genome-wide binding of several TFs in cells derived from WT and DKO mice and find that loss of HMGNs alters the binding of TFs to chromatin, especially at enhancers. Our studies provide insights into molecular mechanisms whereby ubiquitous, non-sequence specific architectural chromatin binding proteins are recruited to regulatory chromatin to modulate transcription factor accessibility and fine tune cell-type specific gene expression programs.

## RESULTS

### HMGN1 and HMGN2 localize to H3K27ac nucleosomes

Analyses of the genome-wide distribution of HMGN1 and HMGN2 in mouse embryonic stem cells (ESCs) (Fig. 1A,B), resting B cells (rBs) (Supplemental Fig. S1A) and embryonic fibroblasts (MEFs) (Supplemental Fig. S1B) reveal that both HMGN1 and HMGN2 variants colocalize with H3K27ac, a histone modification that marks enhancers and promoters (Strahl and Allis 2000; Heintzman et al. 2009; Hnisz et al. 2013), but not with H3K27me3, an epigenetic mark of silent chromatin. Furthermore, in all 3 cell-types, signal but shows no correlation with the H3K27me3 signal levels (Fig.1B, S1A,B). The positive correlation between both HMGN1 and HMGN2 and the H3K27ac signal intensities is especially obvious in the super-enhancer regions of the 3 cell-types examined (Fig. 1C, S1C,D) and it’s slope is noticeably steeper than seen with other epigenetic marks of active chromatin such as H3K9ac, H3K4me1, H3K64ac and H3K122ac (Fig.1D, S1E,F), reflecting the relative high abundance of H3K27ac at super-enhancer regions. The high co-occupancy of HMGN with H3K27ac prompted us to investigate the mechanism whereby HMGNs, which bind to nucleosomes without specificity to DNA sequence are targeted to the H3K27ac nucleosomes.

**Figure 1.**
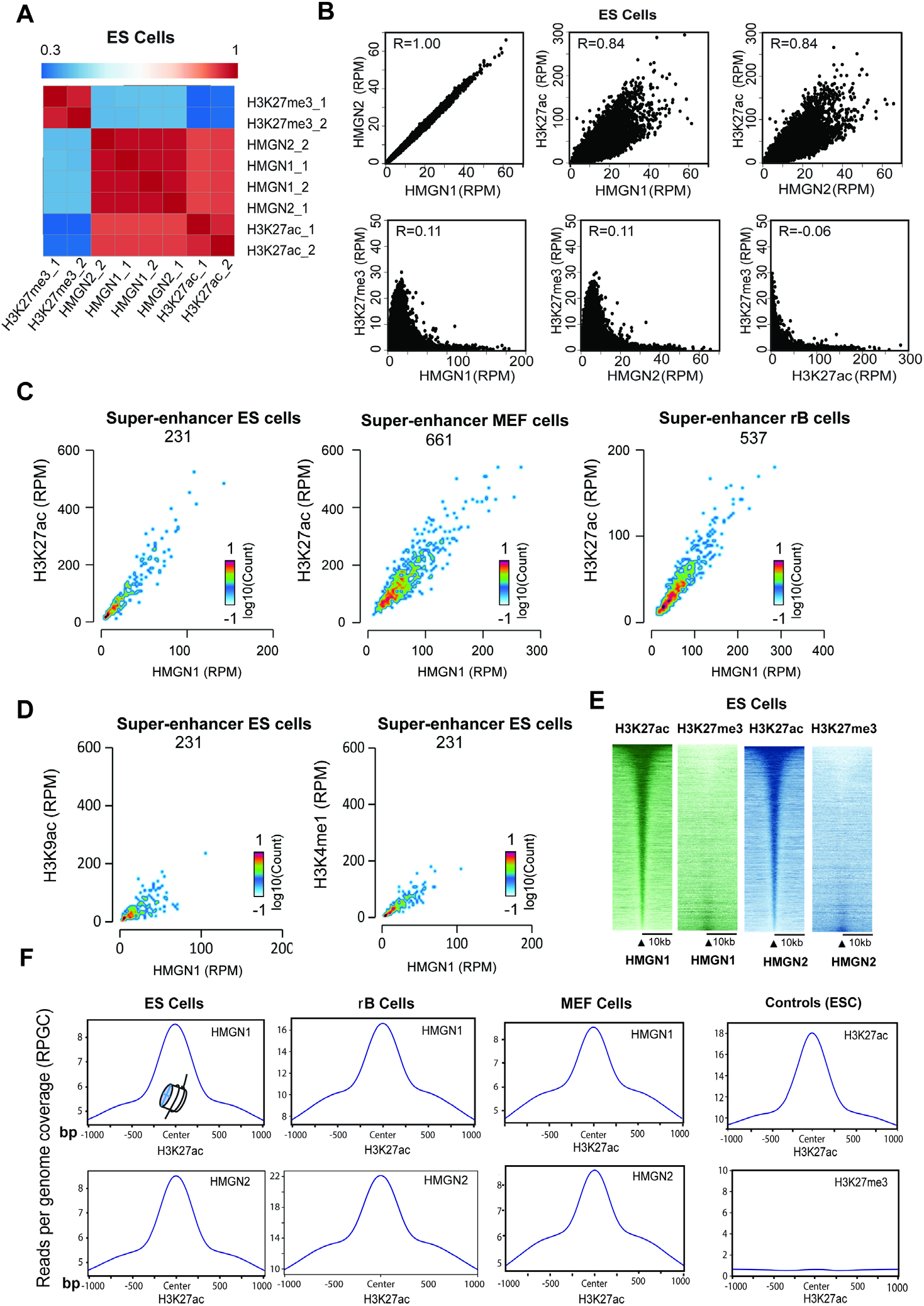
HMGN1 and HMGN2 localize to H3H27ac nucleosomes. **(A)** Correlation heat map showing preferential localization of both HMGN1 and HMGN2 to chromatin regions containing H3K27ac but not to regions containing H3K27me3. Bin size:5000bp. **(B)** Scatter plot showing direct correlation between HMGN1 and HMGN2 occupancy levels and H3K27ac, but not H3K27me3 in ESCs. **(C)** Scatter plot showing direct correlation between occupancy levels of HMGN1 and H3K27ac at the super enhancers of ESCs, MEFs, and resting B cells. The number of super enhancers in each cell-type is indicated below the cell name. **(D)** Scatter plot showing the correlation between occupancy levels of HMGN1 and either H3K9ac or H3K4me1at ESCs super enhancers **(E)** Heat map showing localization of the H3K27ac and H3K27me3 signal at chromatin sites containing either HMGN1 or HMGN2. **(F)** High resolution profile plots showing genome-wide co-localization of HMGNs and H3K27ac in several cell-types. ES: embryonic stem cells; MEF: mouse embryonic fibroblasts; rB: resting B cells.

Throughout the genomes of the three cell-types, the H3K27ac signal, but not the H3K27me3 signal centers at the chromatin loci containing either HMGN1 or HMGN2 (Fig 1E, S1G). Significantly, high resolution analyses of the colocalization plots reveal that the location of both HMGN1 and HMGN2 centers narrowly on the location of the H3K27ac signal, with the same precision as the H3K27ac signal itself (Fig. 1F). The overlap between the H3K27ac and the HMGN chromatin occupancy signals suggest that HMGNs localize to the H3K27ac nucleosomes.

In sum, the ChIP seq data indicate that both HMGN1 and HMGN2 localize to nucleosomes containing the H3K27ac modification, one of the most abundant epigenetic marks of active chromatin regulatory sites. Given that H3K27ac marks cell-type specific chromatin regulatory sites (Heintzman et al., 2009; Hnisz et al., 2013), these findings provide insights into the molecular mechanism whereby HMGNs affect cell-type specific gene expression.

### Mechanism targeting HMGNs to H3K27ac nucleosomes

To further verify that HMGNs preferentially bind to H3K27ac nucleosomes and explore the mechanisms that targets these proteins to the acetylated nucleosomes, we prepared chromatin from purified MEF nuclei, stripped the chromatin binding proteins by salt extraction, digested the salt-extracted chromatin with micrococcal nuclease, fractionated the digest on a sucrose gradient, and isolated chromatin fragments containing either only mononucleosomes (MN) or oligonucleosomes (ON); a mixture of tri-penta nucleosomes (Fig. 2A). To the ON fraction we added purified HMGN1 or HMGN2 at a ratio of one molecule of HMGN per 25 nucleosomes and immunopurified the ON fraction containing bound HMGN, with antibody to HMGNs. We purified the H3 histone fraction from the input and from the immunopurified ON by HPLC and performed dot-blot western analysis with antibodies to either H3K27ac, H3, H3K27me3 or H3K9ac.

**Figure 2.**
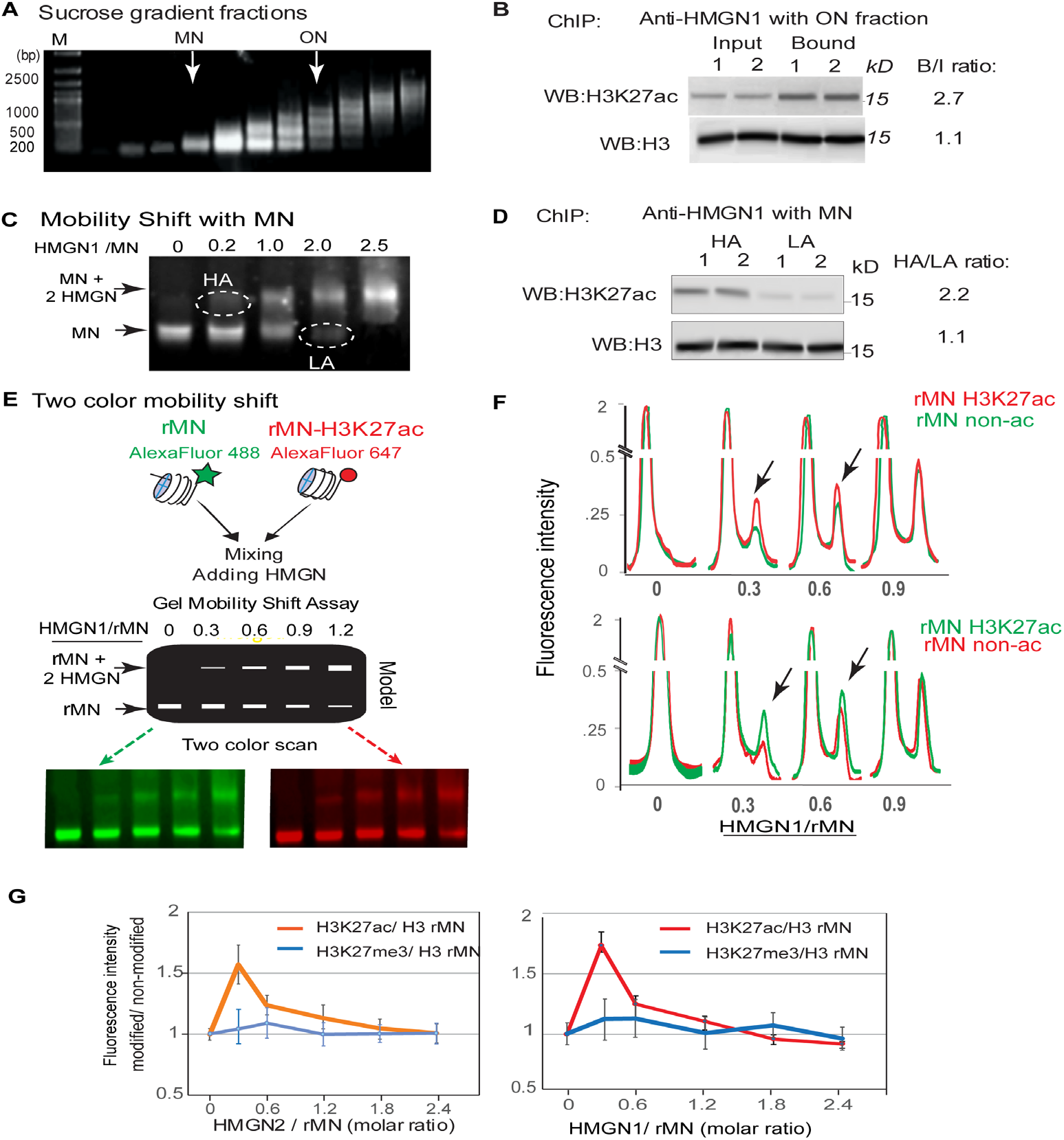
Preferential binding of HMGN to chromatin particles containing H3K27ac. **(A)** Agarose gel showing sucrose gradient fractionated salt stripped MEF chromatin particles. MN: mononucleosomes, ON: Oligonucleosome (mostly tri-penta nucleosomes). **(B)** Western analysis of total ON (Input) and HMGN1 immunoprecipitated ON (bound) **(C)** Gel mobility shift -assay. Purified HMGN1 was added to salt stripped chicken erythrocyte MNs at the ratio indicated on top of each column. The MNs shifted at low HMGN1:MN were designated as high affinity (HA) while the MNs not shifted at high HMGN:MN were designated as low affinity (LA). **(D)**. Western analysis of HA and LA mononucleosomes (MN) **(E)** Two color gel mobility shift assays of recombinant mononucleosomes (rMN). A mix of equal amounts of fluorescently Alexa 488 labeled rMN (green) and Alexa 647 labeled rMNH3K27ac (red) were incubated with various amounts of HMGN1, the mixture fractionated on native polyacrylamide gels, and the gels scanned to visualize and quantify either the red or green fluorescence. Shown is the experimental design and gel images visualized with red or green channels **(F)** Scan of the gels shown in E and of a similar gel in which the fluorescent labels are reversed. Top: Alexa 647 labeled rMNH3K27ac (red) and Alexa 488 labeled rMN (green). Bottom: Alexa 488 labeled rMNH3K27ac and Alexa 647 labeled rMN. Arrows point to preferential binding of HMGN to rMNH3K27ac in the shifted nucleosome. **(G)** Quantification of the scans shown in panel F and in Supplemental Fig. S3 for HMGN1 and of similar experiments done with HMGN2. Note that at low ratio of HMGN to nucleosomes, both HMGN1 and HMGN2 preferentially bind to rMNH3K27ac but not to rMNH3K27me3.

These dot-blot analyses reveal that the levels of H3K27ac in ON fraction that bound either HMGN1 or HMGN2 are over 3-fold higher than in the input ON fractions (Fig. 2A), an indication that *in vitro*, HMGNs preferentially interact with chromatin fragments enriched in nucleosomes containing acetylated H3K27, but not with fragments enriched in either H3K27me3 or H3K9ac. Western analysis verified the preferential binding of HMGN1 to H3K27ac nucleosome; when comparing the H3K27ac signal ratio between input and HMGN1 bound nucleosomes we find that the levels of H3K27ac signal in HMGN bound nucleosomes was 2.7 fold higher than in input nucleosomes (Fig. 2B), further indication that HMGNs bind preferentially to nucleosomes containing H3K27ac.

Purified HMGN proteins bind specifically to MNs and form complexes (HMGN:MN) which in a mobility shift assay migrate slower than free MN particles, that do not contain HMGN (Mardian et al. 1980; Crippa et al. 1992) (Fig. 2C). We reasoned that the MN in complexes formed at low HMGN to MN ratio (HA, in Fig. 2C) have a higher affinity for HMGN than MN that did not form complexes even at relatively high HMGN to MN ratios (LA in Fig. 2C). In our experiments, at an HMGN1:MN ratio of 0.2, the HA fraction contained only 9% of the total MNs, while at HMGN:MN ratio of 2.0, the LA fraction contained only 12% of the total MN (Supplemental Fig. S2B). Dot-blot westerns with H3 purified from HA and LA MNs that were complexed with either HMGN1 or HMGN2, indicate that the levels of H3K27ac in the HA complexes are approximately 1.5 times higher than in LA MNs (Supplemental Fig. S2C,D), a further indication that HMGN proteins preferentially bind to MNs containing H3K27ac. Similar experiments using antibodies specific to either H3K27me3 or to H3K9ac did not show enrichment of modified MNs in the HA fraction (Supplemental Fig. S2C,D). In addition, Western analysis of the purified HA and LA mononucleosomes, using antibodies to either H3K27ac or H3 show that the levels of H3K27ac in the HA fraction was 2.0-fold higher than in the LA fraction (Fig. 2D), further evidence that HMGN1 preferentially bind to nucleosomes containing H3K27ac.

Taken together, the results show that HMGNs preferentially bind to nucleosomes and chromatin fragments containing H3K27ac but show no preference for nucleosomes containing other marks of active chromatin such H3K9ac and H3K4me1, or for nucleosomes containing H3K27me3, a mark of transcriptionally silent, compact chromatin.

Native MNs may contain more than one histone modification (Voigt et al. 2012). To test whether the H3K27ac modification, by itself, is enough to preferentially target HMGN to acetylated MNs, we used commercial recombinant MN (rMN) that were either devoid of any modification or contained only the H3K27ac modification (rMNH3K27ac). We end labeled the DNA in rMN with AlexaFluor 488 (green) and the DNA in rMNH3K27ac with AlexaFluor 647 (red) respectively, mixed equal amounts of the red and green fluorescent labeled particles, performed mobility shift assays at increasing molar ratio of HMGN to the fluorescently labeled rMNs, and scanned the gels to visualize either the green or red signal(Fig. 2E). To exclude possible effects of the fluorescent label, we reversed the label and repeated the mobility shift assays with green-labeled rMN and red-labeled rMNH3K27ac. Quantification of fluorescence scans of the gels (Fig. 2F) indicates that at low HMGN to nucleosome ratio the slower moving fraction, which contains HMGN bound nucleosomes (rMN+2HMGN in Fig 2E), was enriched in rMNH3K27ac (Fig. 2 F). As an additional control, we performed the same type of experiments with rMN and rMNH3K27me3 that were labeled with the same fluorochromes. Scans of these gels indicate that HMGNs do not show preferential binding to the H3K37me3 recombinant nucleosomes (Supplemental Fig. S3A,B). Quantitative analyses of the scans of the two-color mobility shift assays show that the ratio of the fluorescence intensity of rMNH3K27ac to rMN was as high as 1.7 while that of H3K27me3/rMN was close to 1.0 (Fig 2G).

In sum, ChIP-Western analysis with purified ONs, or with HMGN-MN complexes purified by gel mobility, and two-color mobility shift analysis with rMN and rMNH3K27ac, all indicate that HMGNs preferentially bind to MNs containing H3K27ac (Supplemental Table 1).

Next, we tested whether the preferential binding of HMGN to MNH3K27ac depends on specific regions present in HMGN proteins or on specific properties of the acetylated MNs. The preferential binding of HMGNs to MNH3K27ac does not solely depend on a direct interaction of the protein with the acetylated H3K27 residue since two color mobility shift assays indicate that the preferential binding of HMGN to MNH3K27ac is maintained even in the presence of 1000 fold molar excess of a competing acetylated peptide (KAARK(27ac)SAPATGG) spanning the H3K27ac residue (Supplemental Fig. S4A). HMGN proteins contain 3 functional domains: a bipartite nuclear localization signal, a highly conserved nucleosome binding domain (NBD) and a regulatory domain which is located in the C-terminal of the protein (Supplemental Fig. S4B) and has been shown to specifically interact with the N-terminal region of histone H3 (Trieschmann et al. 1998; Postnikov and Bustin 2010). It has been demonstrated that HMGN deletion mutants lacking the C-terminal regulatory domain bind to nucleosomes, but mutations in the NBD abolish the binding of HMGN to MN regardless of acetylation status (Prymakowska-Bosak et al. 2001). The HMGN regulatory domain is not directly involved in the recognition of the H3K27ac modified nucleosome since two-color mobility shift assays indicate that a deletion mutant of HMGN1 lacking the C-terminal regulatory domain still preferentially binds to rMNH3K27ac (Supplemental Fig S4B, right). Thus, HMGNs do not contain domains that specifically recognize the acetylated H3K27 residue.

The unstructured N-terminal of histone H3 has been shown to interact with nucleosome linker DNA, and HMGNs have been shown to modify the interaction of the H3 tail with the linker DNA (Murphy et al. 2017). To test whether the linker DNA plays a role in the preferential binding of HMGN to H3K27ac nucleosomes, we digested the 165bp rMN particle with micrococcal nuclease to generate the linker-less 147bp core particles (rCP) (Supplemental Fig. S4C). Two-color mobility shift assays reveal that removal of the DNA linker region abolished the preferential binding of HMGN to rMNH3K27ac particles (Supplemental Fig. S4C, right panel). Thus, even though HMGN does not bind directly to the linker DNA, its presence enhances the preferential binding of HMGN to the acetylated nucleosomes.

In sum, the acetylated H3K27 residue is not the major HMGN1 binding site, and HMGN proteins do not contain specific domains that lead to preferential binding of HMGN to acetylated MNs. Thus, the unique properties of the MNH3K27ac, as compared to non-acetylated MNs, are the major facilitators of the preferential binding of HMGN to the modified MNs.

### HMGNs modulate the levels of H3K27 modifications and histone H1 binding at chromatin regulatory sites

Given the preferential binding of HMGNs to MNH3K27ac, we tested whether they affect the levels of epigenetic modifications at H3K27 residues and performed ChIP analyses with MEFs derived either from WT or from double knock-out mice lacking both HMGN1 and HMGN2 (DKO mice) (Deng et al. 2015). Box plot analysis of these ChIP show decreased levels of H3K27ac, but increased levels of H3K27me3 at both enhancers and promoters of the cells derived from mice lacking HMGNs (Fig. 3A). In agreement, the MA plot show numerous sites, at both enhancers and promoters, where the differences between WT and DKO cells in H3K27ac levels are statistically significant (Fig. 3B, top), and aggregate plots show an increase the global distribution of the H3K27ac only at enhancer regions, where loss of HMGNs leads to a detectable decrease in the H3K27ac occupancy levels (Fig. 3C top lane). For H3K27me3, the MA plots show fewer statistically significant differences between WT and DKO cells in the modification level at a particular site (Fig. 3B,bottom) but aggregate plots indicate that loss of HMGNs leads to a detectable increase in the overall signal of this modification at both enhancer and promoter regions (Fig. 3C, middle lane). Thus, loss of HMGNs decreased the level of H3K27ac but increased the levels of H3K27me3, an indication that HMGNs affect the levels of epigenetic marks at chromatin regulatory sites.

**Figure 3.**
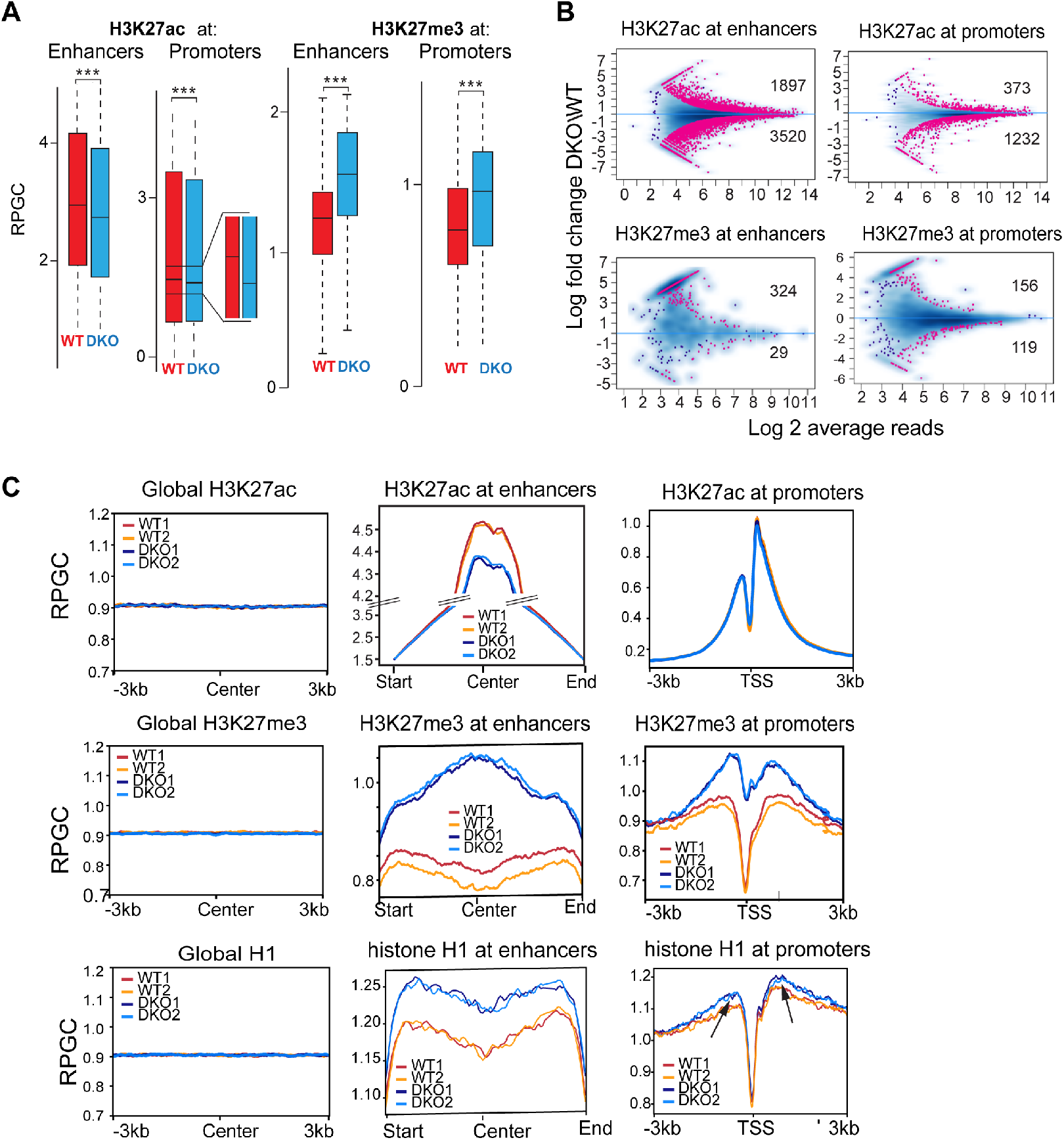
Loss of HMGN alters H3K27modifications and H1 occupancy at chromatin regulatory sites of MEFs. **(A)** Box plot showing decreased H3K27ac but increased H3K27me3 levels at enhancers and promoters of DKO mice. **(B)** MA plots showing differences between WT and DKO cells in H3K27ac or H3K27me3 levels at enhancers and promoters. Statistically significant differences (FDR<0.05) are shown in red. Blue dots and blue density cloud represents all points corresponding to the non-changing regions. **(C)** Aggregate plots showing distribution of the average H3K27ac, H3K27me3 and histone H1 levels. Left: throughout the genome (regulatory sites subtracted) ; center: at enhancers, right: at promoters, in WT and DKO MEFs. Arrows point to promoter regions where H1 occupancy differs between WT and DKO cells. In all panels “Center” indicates a location of the middle point of each 6kbp bin. All ChIP analyses from two biological replicates.

H3K27me3 levels correlate directly with the chromatin occupancy of histone H1 (Kim et al. 2015b; Fyodorov et al. 2018), an abundant protein family that binds dynamically to nucleosomes (Catez et al. 2006) and stabilizes chromatin compaction (Fyodorov et al. 2018). Previous ChIP analyses indicated that H1 is evenly distributed throughout the nucleus but shows low chromatin occupancy at TSS (Cao et al. 2013; Teif et al. 2020). Our ChIP analyses do not show statistically significant differences between WT and DKO in H1 occupancy at a particular site; however, the aggregate plots indicate a marginally higher occupancy of H1 at enhancers and promoters in DKO, as compared to WT regulatory regions (Fig. 3C, bottom lane). Thus, the presence of HMGN lowers the H1 chromatin occupancy at regulatory sites, a finding that agrees with several previous analyses showing that HMGNs destabilize the binding of H1 to nucleosomes (Rochman et al. 2009; Murphy et al. 2017) and reduce its chromatin residence time (Catez et al. 2002).

At transcription start sites, the occupancy levels of both H3K27ac and HMGN show a direct correlation with gene expression levels (Karmodiya et al. 2012; He et al. 2018). We ranked the genes into 5 categories according to their expression levels and noted that indeed, the most highly expressed genes show the highest level of H3K27ac modification and the highest level of HMGN occupancy (Supplemental Fig. S5A). Comparative analysis of WT and DKO cells show that loss of HMGNs increases both the H3K27me3 levels and H1 occupancy at the TSS of highly expressed genes to a markedly larger degree than at low expressing genes (Supplemental Fig. S5B,C), a finding that is in agreement with the preferential occupancy of HMGNs on H3K27ac nucleosomes. In sum, loss of HMGN leads to an altered profile of the epigenetic marks at H3K27 residues, resulting in a relative increase in H3K27me3 levels and H1 occupancy at enhancers and promoters.

### HMGNs affect the binding of regulatory factors to chromatin

Enhancers and promoters serve as major binding sites for transcription factors (TFs). The preferential location of HMGNs at these sites and the epigenetic changes observed in DKO MEFs, together with the known effect of HMGNs on cellular transcription profiles (Kugler et al. 2013) and on the stability of cell identity (He et al. 2018), raises the possibility that HMGNs affect the interaction of TFs with chromatin regulatory sites. Therefore we examined the effect of HMGNs on the chromatin occupancy of the acetyltransferase p300 and of the bromodomain-containing protein Brd3, a “writer” and “reader” of H3K27ac. RNA-seq and western blot analyses of extracts prepared from WT and DKO MEFs, indicate that loss of HMGN did not affect the transcript and protein levels of either p300 (Fig.4A) or Brd3 (Fig.4E). Yet, ChIP analyses reveal that loss of HMGNs leads to a marked decrease in the chromatin occupancy of both p300 (Fig.4 B,C,D,I) and of Brd3 (Fig.4 F,G,H,I) throughout the MEF genome and at both enhancers and promoters. At enhancers and promoters, the number of significantly decreased P300 binding sites (869 and 427, respectively) was 20-fold higher than the sites that show increased occupancy (Fig. 4B,C). The chromatin binding of Brd3 was affected to a larger degree; at enhancers loss of HMGNs decreased significantly Brd3 binding at 5862 sites but increased the Brd3 binding at only 64 sites (Fig. 4G). Similar effects are seen at MEF promoters where the loss of HMGN decreased Brd3 binding at 5090 sites (Fig. 4G). In agreement, aggregate plots show decrease chromatin occupancy of P300 and Brd3 at both enhancers and promoters (Fig.4 D,H).

**Figure 4.**
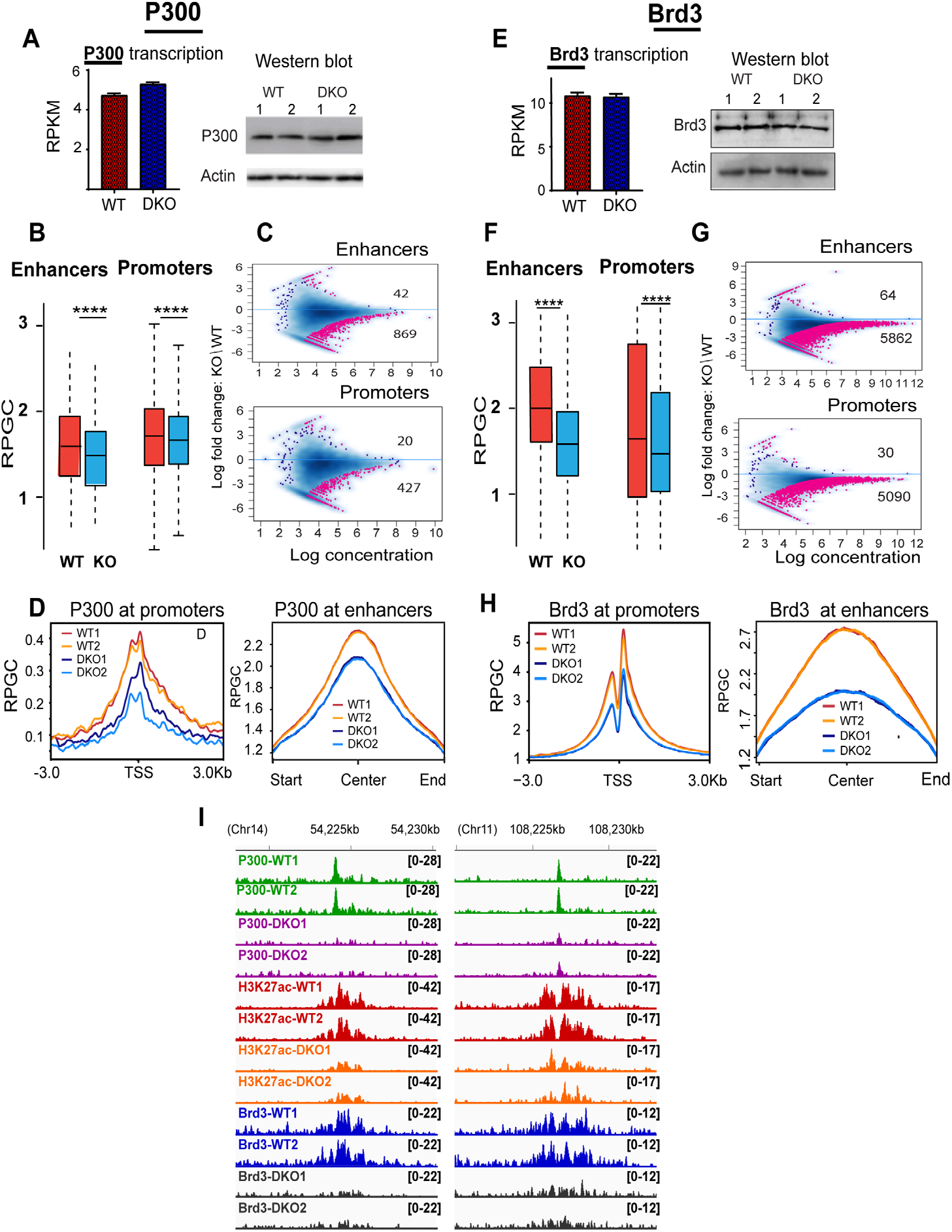
Decreased p300 and Brd3 chromatin binding in DKO MEFs. **(A)** Equal P300 expression in WT and DKO MEFs. **(B)** Box plots showing decrease P300 chromatin occupancy at enhancers and promoters of DKO cells. **(C)** MA plots showing differences in P300 chromatin binding between DKO and WT cells. Sites showing statistically significant differences (FDR<0.05) are in red. Blue dots and blue density cloud represents all points corresponding to the non-changing regions. Note that most altered sites show decrease binding in DKO cells. **(D)** Profile plots showing decreased P300 occupancy at promoters and enhancers of DKO cells. **(E)** Equal Brd3 expression in WT and DKO MEFs. **(F)** Box plots showing decrease Brd3 chromatin occupancy at enhancer and promoters of DKO MEFs. **(G)** MA plot showing differences in Brd3 chromatin binding between DKO and WT cells. Sites showing significant differences (FDR<0.05) are in red. Blue dots and blue density cloud represents all points corresponding to the non-changing regions. **(H)** Profile plots showing decreased Brd3 occupancy at promoters and enhancers of DKO cells. **(I)** IGV tracks showing reduced H3K27ac levels, and reduced P300 or Brd3 chromatin occupancy in DKO cells. All ChIP analyses from two biological replicates.

Likewise, the expression levels of CEBPB, a TF that binds to chromatin with DNA sequence specificity (Koldin et al. 1995), were not affected by the loss of HMGN (Fig. 5A), but the CEBPB chromatin occupancy in DKO MEFs is noticeably diminished throughout the genome (Fig. 5B) and at both promoters and enhancers (Fig 5C). Of the 19,392 CEBPB sites detected in WT cells, 10,865 and 8257 sites were lost and retained, respectively in DKO MEFs (Fig. 5D). The top CEBPB binding sequence motif in both WT and DKO cells corresponds to the canonical CEBPB binding motif (Fig. 5E), an indication that HMGN does not affect the DNA binding sequence specificity of CEBPB. For both lost and retained sites, the CEBPB ChIP seq peaks align narrowly on the center of the CEBPB binding motif, (Fig. 5F, top two lines); however, a search for TF DNA-binding sequence motifs uniquely present only in retained or only in lost CEBPB sites show differences between these sites. The retained CEBPB sites are flanked by DNA binding motifs for additional transcription factors such as FOSB, JUN and ATF3 (Fig. 5F, third line), while the lost CEBPB sites are not surrounded by known transcription factor binding motifs (Fig. 5F bottom line). In addition, the H3K27ac levels at retained sites were higher than at the lost sites (Fig. 5G). In WT MEFs, 64% of CEBPB binding sites localized to chromatin regions showing H3K27ac reads; 86% of these also showed HMGN1 and HMGN2 occupancy. At nonacetylated sites, the co-occupancy of CEBPB with HMGN was only 39% (Fig. 5H, I).

**Figure 5.**
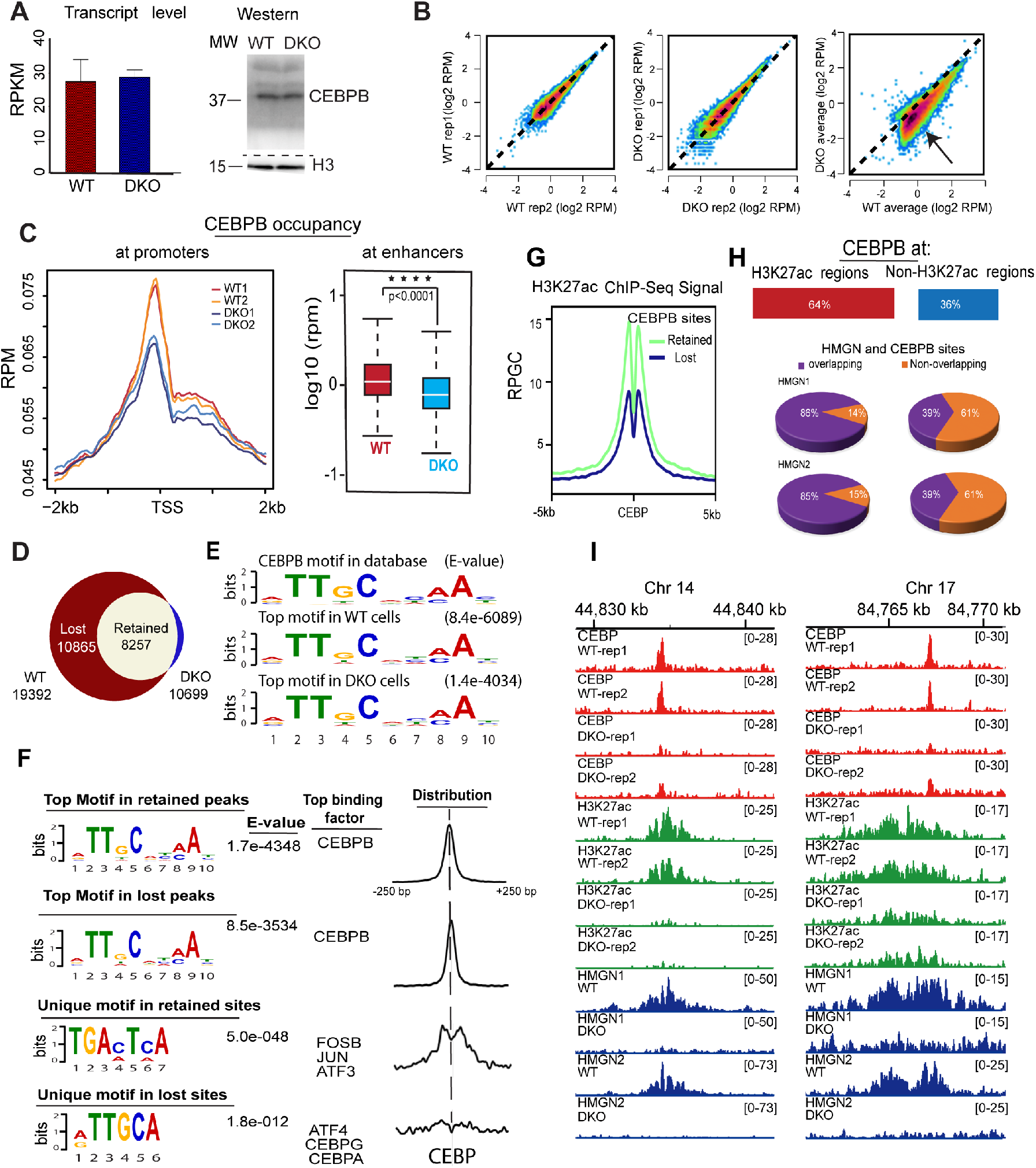
Decreased CEBPB chromatin binding in DKO MEFs. **(A)** Equal levels of CEBPB transcript and protein in WT and DKO MEFs**. (B)** Scatter plot comparing intensities of CEBPB peaks between biological replicates of WT (left), and of DKO cells (center). Right scatter plot shows reduced CEBPB chromatin binding in DKO cells. **(C)** Decreased CEBPB binding at TSS and enhancers of DKO cells. **(D)** Venn diagram showing CEBPB chromatin binding sites in WT and DKO MEFs. **(E)** Top DNA sequence motif underlying the CEBPB binding sites in WT and DKO cells, compared with the CEBPB motif in database. **(F)** Top and unique motifs in retained and lost CEBPB binding sites. Lost CEBP sites are defined as present in WT but not in DKO cells. The diagrams to the right show the location of the DNA binding motifs relative to the center of the CEBPB binding motif. **(G)** H3K27ac levels at lost or retained CEBPB sites in WT cells. **(H)** Overlap between CEBPB, H3K27ac and HMGN occupancy. **(I)** IGV snapshots showing loss of CEBPB binding in DKO cells at regions overlapping with H3K27ac. All ChIP analyses from two biological replicates.

Thus, the binding of CEBPB to chromatin is modulated by, but not exclusively dependent on, the presence of HMGN protein. CEBPB binding sites that contain motifs for additional transcription factors or show high H3K27ac reads are less affected by the loss of HMGNs than sites that show low H3K27ac levels and are not in proximity to additional TFs.

Similar studies show that HMGNs affect the chromatin interactions of TFs known to play a role in the development and function of mouse B cells. ChIP analyses of IKAROS, ETS1, IRF8 and PAX5 in resting B cells isolated from WT and DKO mice show that invariably, loss of HMGN reduced the binding of the TFs to chromatin throughout the genome and at both enhancers and promoters (Supplemental Fig. S6).

In ESCs, loss of HMGN leads to a marked reduction in the chromatin binding of the H3K27ac reader Brd4 (Fig.6, top lane), and to a more moderate loss of chromatin binding of Klf4 (Fig. 6, 2^nd^lane) and CTCF (Fig. 6, 3^rd^ lane), two TFs known to affect global chromatin topology (Di Giammartino et al. 2019). Interestingly, the pluripotency factors NANOG, SOX2 and OCT4, whose nucleosome binding motifs are located in proximity to HMGN binding sites, show higher chromatin occupancy in DKO ESCs than in WT cells (Fig.6, bottom 3 lanes).

**Figure 6.**
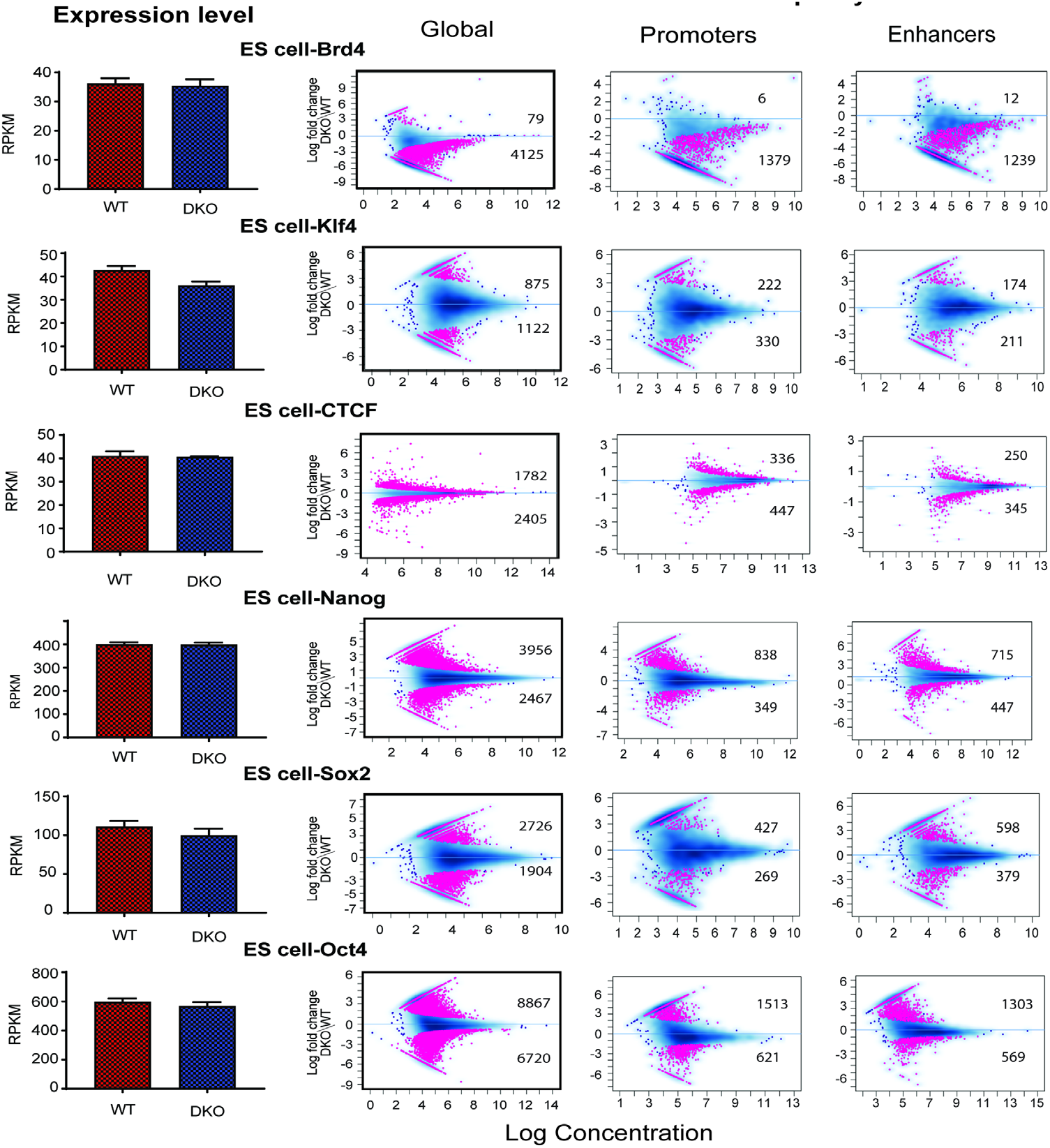
Altered chromatin occupancy of transcription factors in DKO ESCs cells. Bar graphs on left show transcript levels determined by RNA seq analysis. MA plots show differences in TF chromatin binding between DKO and WT cells. Sites showing significant differences (FDR<0.05) are in red; the number of up- and down-regulated sites are indicated in each panel. Blue dots and blue density cloud represent all points corresponding to the non-changing regions All data from 2 biological replicates.

Taken together, ChIP analyses in MEFs, rBs and ESCs consistently show that loss of HMGN alters the interaction of TFs with chromatin; however, the magnitude and type of effect is context-dependent on the exact mode of interaction of a TF with its cognate binding site in chromatin.

## DISCUSSION

H3K27ac is a major epigenetic mark of active chromatin while HMGNs are ubiquitous nuclear proteins that bind dynamically to chromatin without specificity for DNA sequence. Here we identify H3K27ac as an epigenetic signal that facilitates the localization of HMGNs to specific chromatin sites, thereby leading to epigenetic changes that affect the binding of TFs to chromatin. Likely, H3K27ac is not the only epigenetic mark that affects the binding of HMGN to chromatin; additional epigenetic marks may also contribute to the preferential localization of HMGN to regulatory sites. Nevertheless, our findings that H3K27ac-modified nucleosomes help recruit HMGN proteins to chromatin regulatory sites provide novel insights into epigenetic mechanisms that fine-tune cell-type specific transcription and stabilize cell identity.

We find that genome wide, HMGNs colocalize with H3K27ac because they bind preferentially to H3K27 acetylated MNs, as compared to nonacetylated MNs. This preference is seen in experiments containing only purified HMGN and recombinant MNH3K27ac, an indication that the increased binding of HMGN to chromatin regions containing H3K27ac is not dependent on co-factors that facilitate the binding of HMGN to the modified nucleosome, on unique nucleosome spacing, or on the presence of additional histone modifications on the targeted nucleosomes. Thus, the presence of the acetyl residue on H3K27 by itself, is sufficient to preferentially target HMGNs to MNH3K27ac. Considering the mechanisms driving this preference, we note that previous studies show that HMGNs bind to MNs through a conserved nucleosome binding domain that contacts the nucleosome near the nucleosome dyad (Alfonso et al. 1994) and at the nucleosome acidic patch in the H2A.H2B dimer (Kato et al. 2011). The HMGN C-terminal contacts the N-terminal of H3 (Trieschmann et al. 1998) and disrupts its interaction with the linker DNA (Murphy et al. 2017), yet we find that deletion of the HMGN C-terminal domain does not abolish the preferential binding of HMGN to MNH3K27ac. These analyses suggest that an HMGN protein does not contain specific regions that can distinguish between acetylated and non-acetylated MNs. Most likely structural differences between MN and MNH3K27ac lead to the preferential binding of HMGNs to the acetylated MNs.

We identify two major factors that determine the preferential binding of HMGN to MNH3K27ac: the presence of the acetylated H3K27 residue and the presence of linker DNA, yet we find that the HMGNs interaction with the acetylated H3K27 residue is not the major factor determining the preferential binding to acetylated MNs and that HMGNs do bind well to the linker-less 146bp core particle. In considering how acetylation of H3K27 could affect HMGN binding, we note that the flexible and unstructured H3 tail has been shown to interact with both the linker DNA and with the DNA surrounding the histone octamer. The interaction of the H3 tail with nucleosomal DNA can affect nucleosome dynamics (Shaytan et al. 2016; Zhou et al. 2019) and alter the DNA conformations, especially in regions close to the nucleosome dyad axis, a region that has been mapped as a major site where HMGNs bind to the nucleosome (Alfonso et al. 1994; Murphy et al. 2017). Single charge neutralizing modifications such as lysine acetylation can alter the local conformation of the H3 histone tail and its interaction with the DNA (Kim et al. 2015a). Thus, taken together with previous information, our results suggest that acetylation of H3K27 leads to conformational changes in the nucleosome that facilitate binding of HMGN, thereby increasing the time that an HMGN molecule resides at a specific chromatin regulatory site. We note that our findings do not exclude the possibility that additional epigenetic modification affect the binding of HMGN to chromatin.

An important consequence of increased HMGN chromatin residence time is a decrease in the chromatin residence time of the linker histone H1 (Catez et al. 2002), a protein known to promote chromatin compaction (Fyodorov et al. 2018). HMGNs alter the interaction of the histone H1-C terminal with linker DNA (Murphy et al. 2017) and destabilize the interaction of H1 with chromatin. Conversely, a decrease in HMGN levels enhances the interaction of H1 with chromatin, as we previously observed in studies of specific genomic loci (Deng et al. 2017; Zhang et al. 2019) and in this study globally, using cells derived from mice lacking both HMGN1 and HMGN2 (DKO cells). These effects are most obvious at enhancers and promoters, regions which are marked by a high level of MNH3K27ac and high HMGN occupancy. The most highly acetylated promoters, which also show the highest HMGN occupancy in WT cells, show the highest increase in H1 occupancy in DKO cells (Supplemental Fig. S5). Histone H1 facilitates the binding of PRC2-EZH2 to chromatin and stimulate the methylation of H3K27 (Yuan et al. 2012; Fyodorov et al. 2018). In agreement with these studies, DKO cells show increased H3K27me3 levels at enhancers and the highest incremental increase in H3K27me3 levels at the most active promoters, i.e. promoters that in WT cells showed the highest acetylation levels and HMGN occupancy (Supplemental Fig. S5).

Given the preferential binding of HMGN to MNH3K27ac in active chromatin, it could be expected that loss of HMGN would affect the binding of TFs to chromatin. Indeed, we find that loss of HMGNs decreased the chromatin binding of most of the TFs analyzed with the most prominent effects on chromatin regulators that interact directly with H3K27 residue such as p300, Brd3 and Brd4; their chromatin binding was markedly reduced in cell lacking HMGNs. A more detailed analysis of CEBPB binding suggests that HMGNs do not alter the TF DNA binding specificity and that the HMGN effects depend on several additional factors, including the local levels of acetylation and the presence of cofactors that affect the binding of a specific TF to chromatin. Loss of HMGNs do not always reduce the binding of TFs to chromatin. We find that Sox2, Oct4 and Nanog, TFs whose binding sites (Dodonova et al. 2020; Michael et al. 2020) are in proximity to the nucleosomal binding sites of HMGNs (Alfonso et al. 1994), show increased chromatin binding in DKO. Interestingly, the interaction of Sox2 and Oct4 with nucleosomes can lead to detachment of DNA termini from the histone octamer and to distortion in the histone-DNA contacts. The HMGN nucleosome binding sites have been mapped to the major groves flanking the nucleosome dyad axis (Alfonso et al. 1994) and thermal denaturation studies indicate that HMGNs stabilize the structure of MN by minimizing the unraveling of the DNA strands at the end of the particle (Paton et al. 1983; Crippa et al. 1992). Thus, our findings that the chromatin occupancy of SOX2, NANOG, OCT4 is higher in DKO cells agree with the known location of HMGN on the nucleosome and the effects of HMGN on nucleosome stability. Most likely, for these transcription factors, the presence of HMGN hinders access to their specific binding site in the nucleosome and may hamper their ability to unravel the structure of the nucleosome.

Taken together, the results suggest that the effects of HMGNs depend on the exact mode of TF interaction with chromatin. We note that HMGNs are not the major determinant regulating the binding of TFs chromatin. HMGNs modulate and fine-tune rather than absolutely determine their chromatin binding. Nevertheless, changes in HMGN levels have been shown to alter gene expression and destabilize cell identity, most likely due to alterations in the binding of cell-type specific TFs to chromatin (Chronis et al. 2017; He et al. 2018; Mowery et al. 2018; Nanduri et al. 2020).

The H3K27ac mediated recruitment of HMGN to chromatin regulatory regions provides a molecular mechanism for the experimental findings from many laboratories, which repeatedly show that changes in HMGN levels alter cell-type specific gene expression (Nanduri et al. 2020). Although HMGNs bind dynamically to chromatin and constantly move through the entire nucleus, they preferentially localize to cell-type specific super enhancers, chromatin regions that are enriched in MNH3K27ac. The relatively long residence time of HMGNs at these regulatory sites reduces the chromatin residence of H1 and facilitates TFs access to their specific sites, thereby stabilizing cell identity (Chronis et al. 2017; He et al. 2018; Garza-Manero et al. 2019). Changes in HMGN levels can lead to epigenetic changes that affects the binding of TFs to their specific sites (see model Fig. 7), resulting in cell-type specific changes in gene expression that could affect the cellular phenotype. Indeed, mice lacking both HMGN1 and HMGN2 show multiple phenotypes (Deng et al. 2015), reflecting the ubiquitous HMGN expression in all vertebrate cells. In humans, the increase incidents of B cells acute lymphoblastic leukemia seen in Down syndrome was directly attributed to epigenetic changes and altered transcription mediated by increased HMGN1 levels due to the extra copy of *HMGN1,* which is located on human chromosome 21 (Lane et al. 2014; Mowery et al. 2018). Significantly, in both human and mouse cells, overexpression of HMGN1 leads to upregulation of H3K27ac and downregulation of H3K27me3 levels (Cabal-Hierro et al. 2020), further evidence that altered HMGN levels can lead to epigenetic changes that affect the fidelity of cell-type specific gene expression and impact the cellular phenotype.

**Figure 7.**
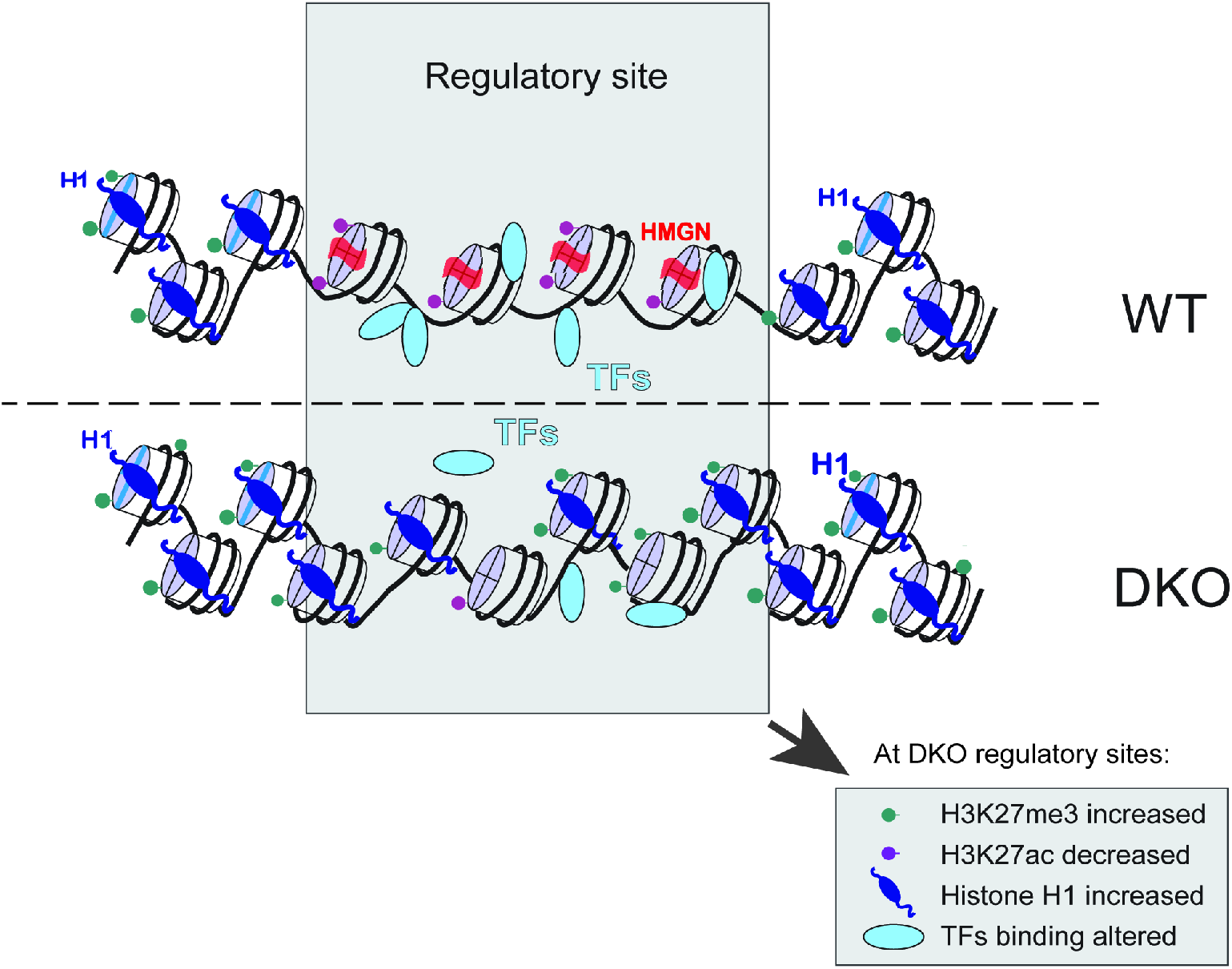
Model of HMGN-mediated epigenetic changes occurring at chromatin regulatory sites. The major changes are indicated in the boxed region at the bottom of the image.

## MATERIALS AND METHODS

### Antibodies, recombinant nucleosomes, peptides and cell lines

Rabbit polyclonal to H1, HMGN1, HMGN2 and H3 were from our laboratory, anti H3K27ac (Abcam#ab4729), anti H3K27me3 (Abcam#ab6002), monoclonal anti H1(Milipore-Sigma #05-457), anti CEBPB (Abcam#ab32358), Anti-Brd3 (Active Motif #61489), Anti-Brd4 (Bethyl Laboratories #A301-985A100), Anti-CEBPB (Abcam #ab32358), Anti-CTCF (EMD Millipore #07-729), Anti-Ets1 (Active Motif #39580), Anti-Ikaros (Active Motif #39355), Anti-Irf8 (Bethyl Laboratories #A304-027A), Anti-Klf4 (Abcam #106629), Anti-Nanog (Active Motif #61419), Anti-Oct4 (Abcam #ab19857), Anti-p300 (Active Motif #61401), Anti-Pax5 (Abcam #183575), Anti-Sox2 (Abcam #97959).

The following recombinant mononucleosomes were purchased from Active Motif : unmodified (#81070); H3K27me3 modified (#81834), H3K27ac modified (#81077). Wild type and HMGN DKO mouse embryonic fibroblasts, embryonic stem cell lines (Deng et al. 2013) and resting B cells (Zhang et al.) were as previously described. Peptides Histone H3 (23 - 34) peptide, KAARKSAPATGG and Histone H3K27ac (23 - 34) peptide, KAAR - K(Ac) – SAPATGG were from AnaSpec, Inc.

### ChIP-Western for HMGN1 and HMGN2

Mono and oligo nucleosomes devoid of protein (salt stripped chromatin particles) were prepared from mouse embryonic fibroblasts (MEFs) derived in our laboratory and from chicken erythrocytes (Rockland R401-0050) as described (Ausio et al. 1989; Postnikov and Bustin 1999a). Briefly, purified nuclei were digested by micrococcal nuclease, the chromatin prepared, non-core histone proteins removed by centrifugation in 0.5M NaCl solutions and cation-exchange chromatography, and the salt-stripped chromatin particles loaded on 5-20% sucrose gradient. Fractions containing either mononucleosomes (MN) and oligonucleosomes (ON, containing 2-5 nucleosomes) were pooled and dialyzed against 10mM NaCl, 10 mM Tris-Cl, pH 7.5, 1mM EDTA. For ChIP-Western analysis the dialyzed nucleosomes were mixed with recombinant HMGN1 or HMGN2 (HMGN:histoneH3 = 50:1 molar ratio) and cross-linked with 1% formaldehyde for 10 minutes at room temperature. After quenching with 0.5M glycine the HMGN-nucleosome complexes were immunoprecipitated using ChIP-IT Express kit (Active Motif, Cat. No. 53008) with either anti-HMGN1, anti-HMGN2 antibodies, or normal rabbit IgG as control. The immunoprecipitated nucleosomes and 2% of the input material were de-cross-linked by heating to 95° for 45 min, and histone H3 isolated by HPLC (Agilent Technologies 1200 series) using a Luna 5 μm CN 100Å HPLC column (Phenomenex, Cat. No. 00D-4255-E0). Equal amounts of H3, as determined by OD220, from bound and input material were blotted on Immobilon PVDF membranes (Millipore, Cat. IPVH304F0), using Schleicher & Schuell Minifold Spot-Blot System and the blots subjected to western analyses.

### Fluorescent labelling of mononucleosomes

For in vitro binding studies, commercial recombinant mononucleosomes (rMN), or rMN containing H3K27 modifications (Active Motif) were first end-labelled with aminoallyl dUTP using recombinant Terminal Deoxynucleotidyl Transferase (TdT), and then concentrated by spin-dialysis (Ultracel 30K, Millipore). The dialyzed rMNs were then labeled with either Alexa Fluor 488 (Green-Fluorescent) or Alexa Fluor 647 (Red-Fluorescent) with succinimidyl ester labeling kits (ARES DNA Labeling Kit, Invitrogen, Cat A21665 and A21676). The fluorescent labelled rMN were concentrated by spin-dialysis.

### Two-color Gel Mobility Shift Assays

Mobility shift assays were performed as described (Postnikov and Bustin 1999b) at ionic strength conditions that lead to binding of 2 molecules of HMGN per nucleosome. Binding reactions contained 100 nM of chromatin particles (25 ng/ul) in 10 μl binding buffer (2×TBE, 0.15 mg/ml BSA and 5% Ficoll). HMGN1, HMGN2 proteins were added to chromatin particles to generate the molar ratio to nucleosomes listed in the figure legends. The mixtures were incubated at 4°C for 10 min and loaded onto non-denaturing 5% polyacrylamide gels in 2× TBE (45 mM Tris-Borate, pH 8.3, 1 mM EDTA) and run at 4°C. The gel was scanned with ChemiDoc MP (BioRad) using duplex fluorescence detection mode (for Alexa Fluor 488 and Alexa Fluor 647). The images were analyzed by Image Lab Touch Software and QuantityOne (BioRad). The ratios between non-shifted nucleosomes (i.e. devoid of HMGN) and shifted (HMGN-bound nucleosomes) were calculated for every titration point.

### Chromatin immunoprecipitation, Illumina library construction and sequencing

The ChIP-Seq procedure was performed as recommended by Active Motif (Carlsbad, CA) Instruction Manuals for ChIP-IT High Sensitivity, ChIP-IT Express, and Chromatin IP purification kits. Briefly, about 10^7^ cultured cells were fixed in medium with 1% formaldehyde (v/v) for 10 min at room temperature on a rocking platform, followed by quenching with 125 mM glycine. Crosslinked cells were washed and incubated in 1 ml Chromatin Prep Buffer containing 1 μl proteinase inhibitor cocktail (PIC) and 1 μl of 100 mM PMSF for 10 min on ice followed by centrifugation at 1250× g for 3 min at 4°C. The pellets were re-suspended in 250 μl ChIP Buffer with 2.5 μl PIC and 2.5 μl 100 mM PMSF and sonicated for 10 cycles with Bioruptor (30 sec on/ 30 sec off). Aliquots of 25 μl of sonicated chromatin were used to generate the input DNA. 5-10 μg of affinity-pure ChIP-grade antibodies) were then added to the rest of the chromatin samples and incubated overnight at 4°C with rotation. Following incubation 30 μl of protein G agarose beads or Magnetic Beads were added to each reaction and the mixtures were further incubated for 3 h or O/N at 4°C. The beads were washed five times with Wash Buffer AM1 (Active Motif). ChIP DNA was eluted in 100 μl Elution Buffer AM4 (Active buffer). Cross-links were reversed at 65°C overnight in the presence of 3 μl of 10% SDS and 5 μl of proteinase K (20 mg/ml). The DNA samples were eluted in 21 μl of elution buffer using MiniElute kit (Qiagen). ChIP-seq library was prepared following the manufacturer’s instructions (Illumina). Briefly, immunoprecipitated and input DNA were blunt ended, ligated to adapters, amplified with PCR and size selected. The ChIP templates were sequenced at 75 bp single read length with Illumina NextSeq 500 system or 101 bp paired end length with HiSeq 2000 by the NIH CCR sequencing facility (for details see data submission file). For the various samples the number of trimmed reads, successfully mapped to the mouse genome, ranged from 12 to 74 million per sample, with an average of 27 million reads with over 80% of trimmed, non-duplicated reads mapped to the genome. Sequence reads were aligned to the Build 37 assembly of the National Center for Biotechnology Information mouse genome data (NCBI37/mm9). Super-enhancers and enhancers were identified as described before (21). Data for the following histone marks were downloaded from GEO archive (accession numbers are indicated in brackets) and processed in the same way as described in the methods section: ES cells - H3K9ac (GSM2417092), H3K4me1 (GSM2629668) and H3K122ac (SRR3144856); MEF cells - H3K9ac (GSM1979773) and H3K4me1 (GSM3272827).

### Quantification and Statistical Analysis

Chromatin binding peaks were initially selected for analysis based on a q-value cutoff 0.01 for broad and 0.05 for narrow regions as reported by the MACS2 peak-calling algorithm (broad or narrow). Peaks were identified as significantly differentially bound using the default threshold of FDR < 0.05. Differentially expressed genes between WT and DKO cells were binned according to the average expression of 3 WT and DKO samples, with 2794 genes assigned to each group. Statistical analyses were performed within the R (ver. 3.6) computing environment and visualized with IGV. Details of statistical analyses can be found in figure legends.

Raw data for H3K27me3 histone modification in B cells [GEO series accession: GSM2184272] were downloaded from NCBI SRA archive and processed in a way identical to other samples. All other ChIP data was generated in our laboratory. Adapter trimming was performed using CutAdapt v.1.16. Sequence quality before and after trimming was checked with FastQC 0.11.5 tool. Sequences were checked for contamination with Kraken v1.1 (https://doi.org/10.1186/gb-2014-15-3-r46) and FastQscreen v.0.9.3 (https://www.bioinformatics.babraham.ac.uk/projects/fastq_screen) applications. Reads were mapped to UCSC mm9 reference genome using BWA 07.17 (https://doi.org/10.1093/bioinformatics/btp324). Duplicate reads were marked with Picard v. 2.17.11. Reads mapped to blacklisted regions (https://doi.org/10.1038/nature11247) were removed from further analysis. For all samples, peak detection of enriched binding regions was performed using either MACS2 v. 2.1.1.20160309 with the default settings, or SICER v. 1.1 with the following parameters: window size 300, gap size 600, FDR < 1e-2, effective genome size 0.75. BigWig files were used for visualization. Correlation heatmaps showing sample relations, scatterplots and peak profiles were generated using DeepTools v. 3.0.1 toolset. Differential binding sites were identified using DiffBind package. To calculate Pearson or Spearman correlation coefficients between different histone mark profiles, the genome was split into bins (bin size =5000 bp) followed by counting numbers of aligned reads within each bin for each sample and calculating a correlation between these sets.

RNA expression levels were determined by RNA-seq as described (Deng et al. 2015; Zhang et al. 2016; Deng et al. 2017). SuperEnhancer Regions were downloaded from SuperDB Superenhancer Database as described (He et al. 2018).

## Data Accessibility

All sequencing data generated as a part of this study are deposited to NCBI and are available with the accession number: GSE156697.

## Competing Interest Statement

The authors declare no competing interests.

## Acknowledgments

This project was funded by the Center for Cancer Research, Intramural Research Program of the National Cancer Institute, NIH and by Contract No. HHSN261200800001E from the National Cancer Institute. The content of this publication does not necessarily reflect the views or policies of the Department of Health and Human Services, nor does mention of trade names, commercial products, or organizations imply endorsement by the U.S. Government. We thank Dr. Ravikanth Nanduri and Bing He for comments on this manuscript.

## Author contributions

SZ and MB conceived and designed the study. SZ, YP, TF, and TD performed experiments and analyzed data. AL and SZ performed bioinformatic analyses. SZ and MB wrote the manuscript, MB coordinated the project. All authors approved the final version of the manuscript. The authors declare no competing interest

---

**Table S1.** The levels of histone H3 modifications in HMGN-bound nucleosomes.

| Substrate | Method | Parameter measured | H3 Modified/H3 |  |  |  |
| --- | --- | --- | --- | --- | --- | --- |
|  |  |  | H3 | H3K27<sup>ac</sup> | H3K9<sup>ac</sup> | H3K27me<sup>3</sup> |
| ON MEF | ChIP(anti-HMGN1)-WB | B/I | 1 | 2.7 |  |  |
| MN RBC | GMSA (HMGN1) –WB | HA/LA | 1 | 2.2 |  |  |
| MN RBC | GMSA (HMGN1) – dWB | HA/LA | 1 | 2.3 | 1.1 | 0.9 |
| MN RBC | GMSA (HMGN2) - dWB | HA/LA | 1 | 1.6 | 1 | 0.9 |
| ON MEF | ChIP(anti-HMGN1)-dWB | B/I | 1 | 3.1 | 0.8 | 0.9 |
| ON MEF | ChIP(anti-HMGN2)-dWB | B/I | 1 | 3.3 | 1 | 0.9 |
| rMN | dual-color GMSA (HMGN1) | red/green | 1 | 1.7 |  | 1.1 |
| rMN | dual-color GMSA (HMGN2) | red/green | 1 | 1.6 |  | 1.1 |

**Supplemental Figure S1.**
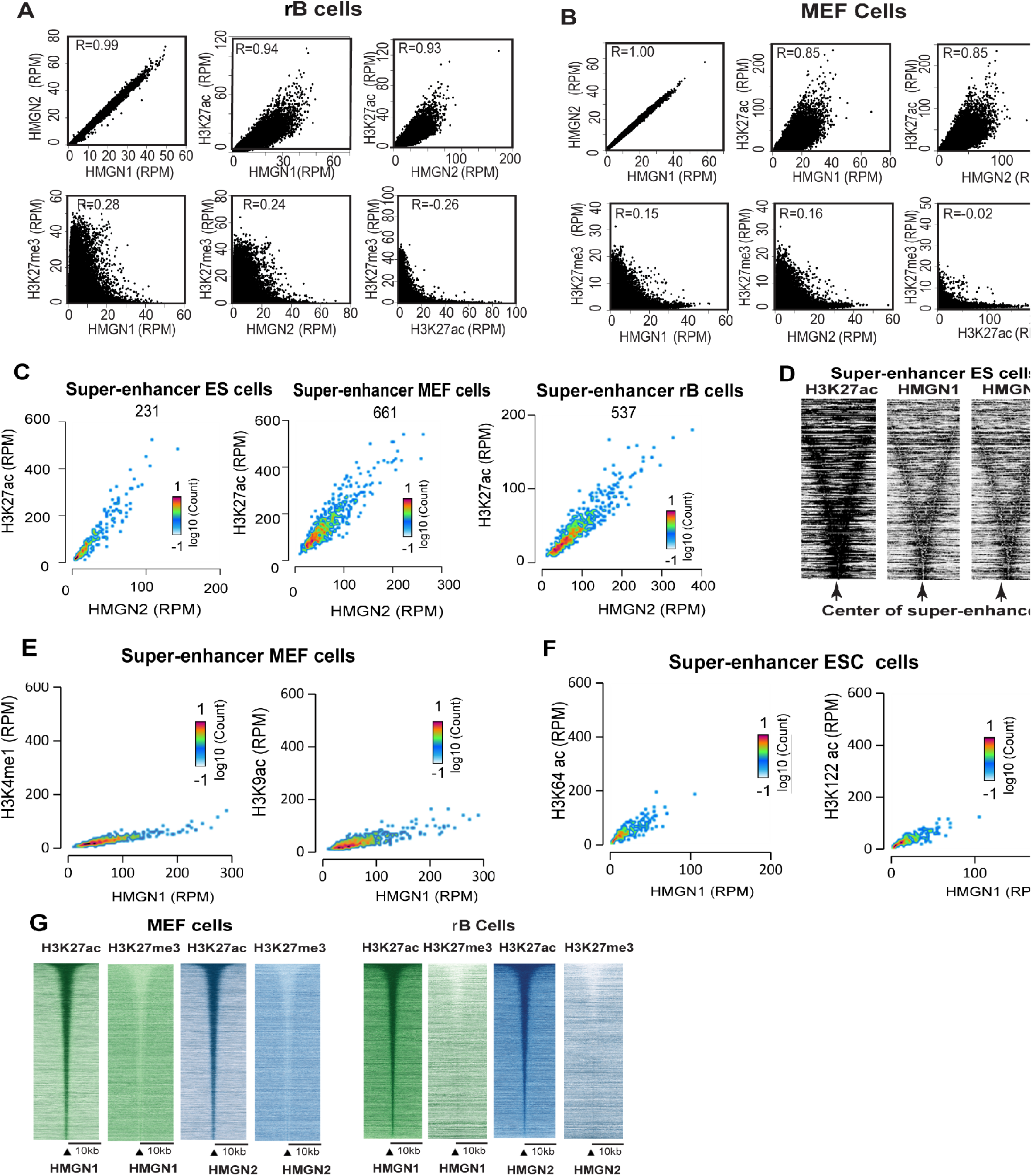
HMGNs localize to H3K27ac nucleosomes. **(A)**. Scatter plot showing a direct correlation between HMGN occupancy levels and H3K27ac, but not H3K27me3 in rBs (**B)**. Scatter plot showing a direct correlation between HMGN occupancy levels and H3K27ac, but not H3K27me3 in MEFs (**C).** Scatter plot showing direct correlation between HMGN2 and H3K27ac occupancy levels at the specific super-enhancers of ESCs, MEFs and rB cells. The number of super-enhancers in each cell-type is shown. (**D)**. Heat map showing correlation between the occupancy levels of HMGN1, HMGN2 and H3K27ac within the ESCs super-enhancers. (**E).** Scatter plot showing the correlation between HMGN1 occupancy levels and H3K4me1 and H3K9ac at MEF super enhancers. (**F).** Scatter plot showing correlation between HMGN1 occupancy levels and H3K64ac and H3K122ac at mouse ESCs super-enhancers. **(G)** Heat map showing localization of the H3K27ac, but not of the H3K27me3 signal at chromatin sites containing either HMGN1 or HMGN2 in MEFs and rB cells.

**Supplemental Figure S2.**
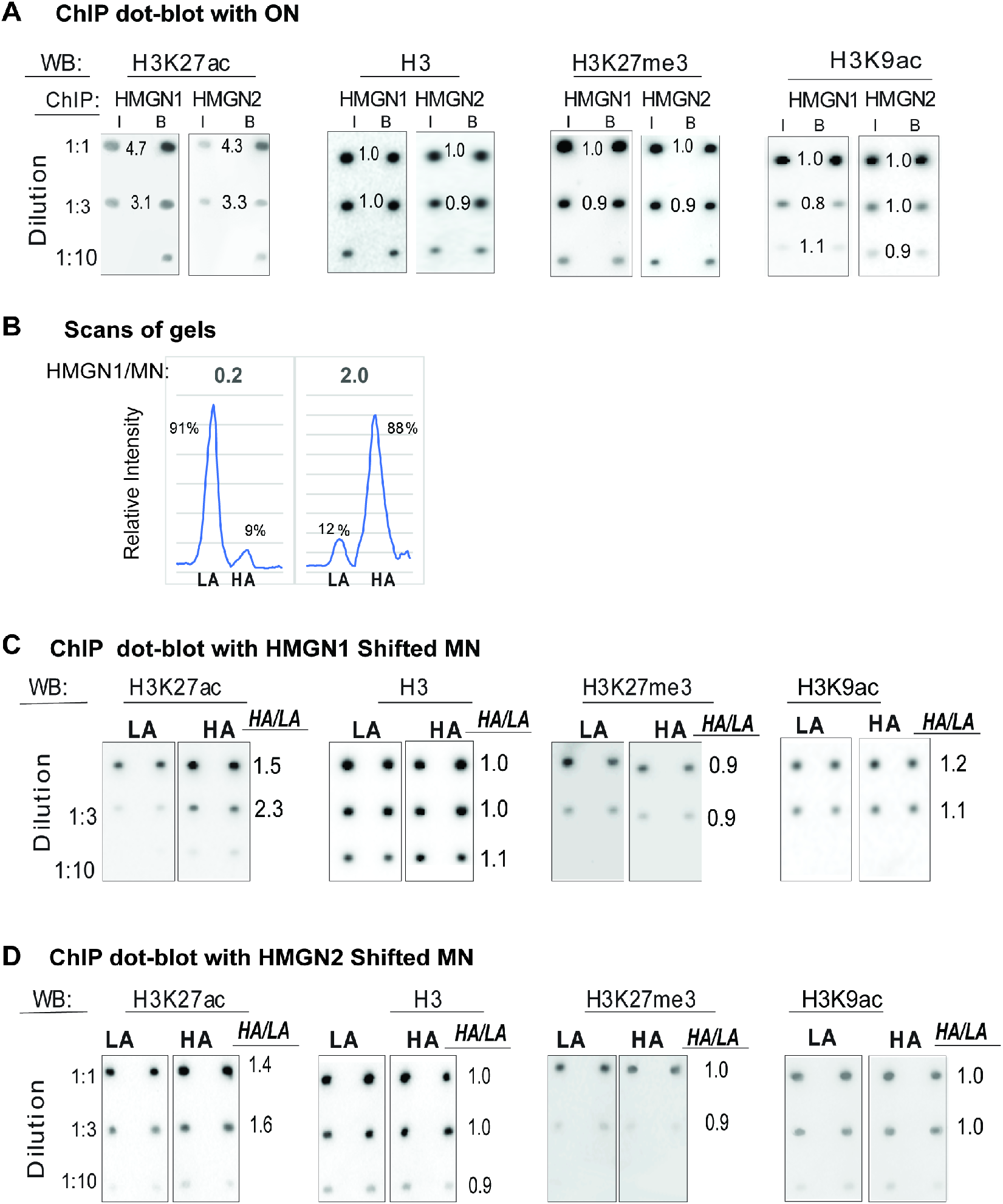
Preferential binding of HMGNs to H3K27ac nucleosomes. **(A)** ChIP dot-blot western with ON. HMGN1 or HMGN2 added to ON fraction (HMGN:histone H3 = 1:50), the complexes immunoprecipitated, the histone H3 from input (I) and HMGN-bound **(B)** ON purified, applied at 3 separate dilutions to PVD membranes, the membranes subjected to Western analysis with the antibodies indicated at top of the panels, and the intensity of the signal quantified. The numbers between the dots indicate the B/I ratio. Note the high B/I ratio for H3K27ac but not for H3K27me3, H3K9ac or H3 which serves as a loading control. Two strips were tested for each experiment. (**B)**. Scans of selected lanes from Fig. 2C showing the percent of nucleosomes designated as either HA or LA. (**C,D)** Dot-blot Western analysis of LA and HA particles. The relative levels of H3K27ac in purified H3 determined as described in panel A except that the mobility shift was done on an HMGN:MN =2. Note the relative high HA/LA ratio for H3K27ac, but not for the three controls. Each strip contains duplicate samples; two strips were tested for each experiment.

**Supplemental Figure S3.**
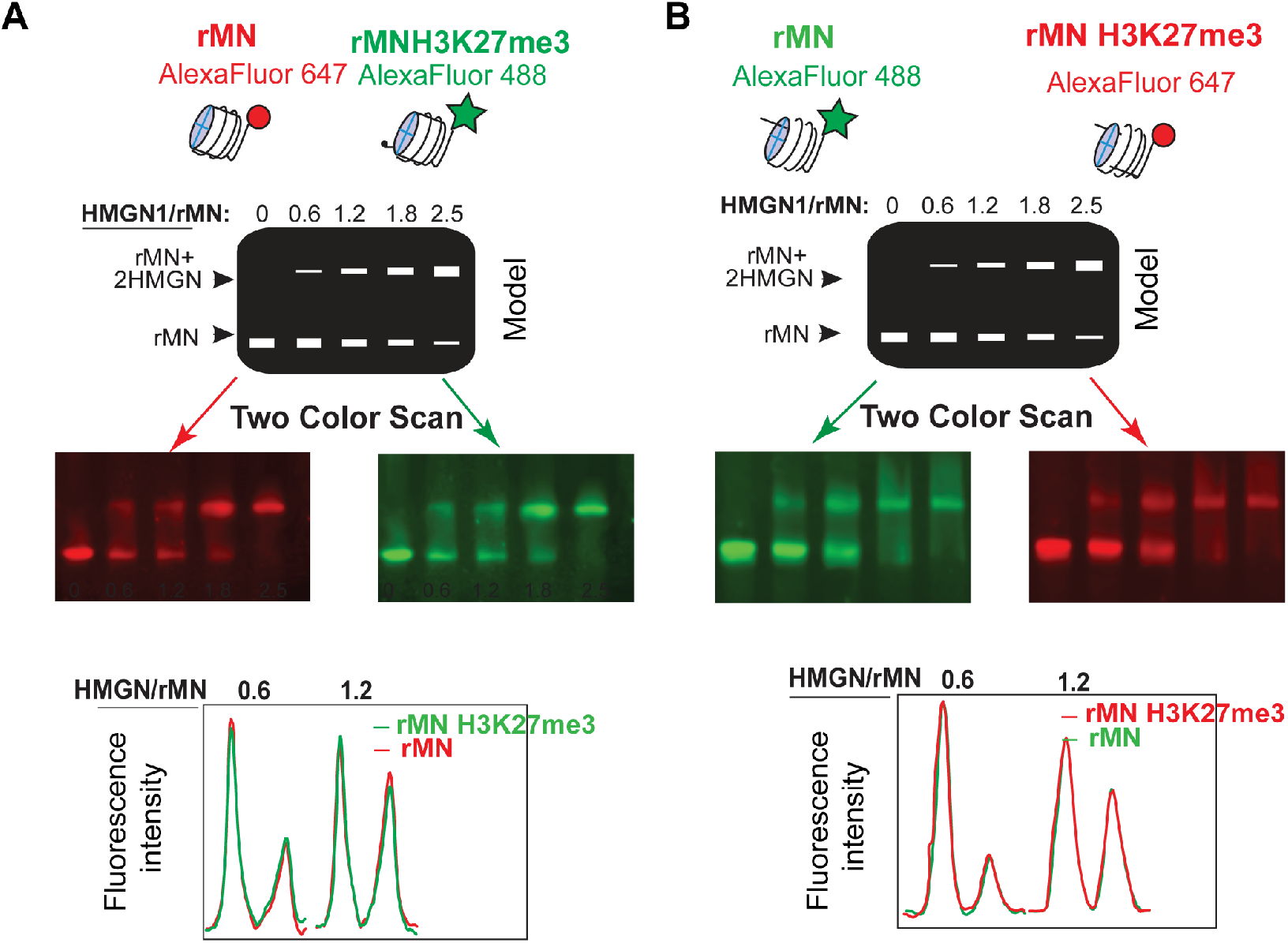
HMGN1 does not bind specifically to the H3K27me3 MNs. **(A)** Two color gel mobility assays of a mixture Alexafluor 647 labeled rMN (red) and Alexafluor 488 labelled rMNH3K27me3 (green). (B) As in (A) with the color label reversed. Scans of selected lanes shown below the gel images. Quantification of the scan shown in Figure 2G.

**Supplemental Figure S4.**
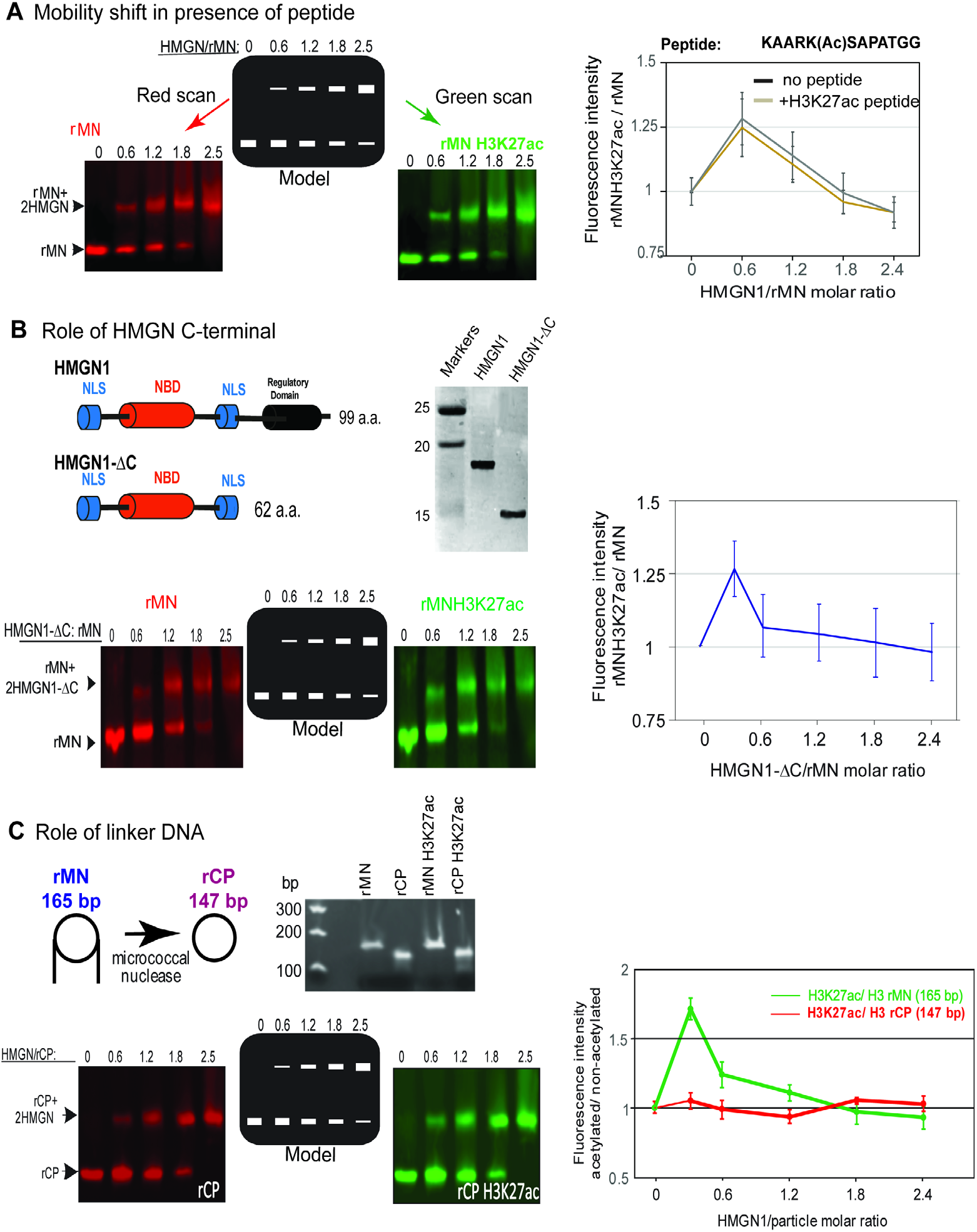
Determinants of preferential binding of HMGN to MNH3K27ac. **(A)** HMGN1 does not bind specifically to the H3K27ac residue. Gel images of two-color mobility shift assays done in the presence of 1000-fold molar excess of a peptide spanning residues 23-34 of H3 in which residue K27 was acetylated. Sequence of the peptide and quantification of the gels is shown on the right. **(B)** The C-terminal domain of HMGN1 is not necessary for preferential binding to rMNH3K27ac. Shown is a schematic diagram of full length and of the C terminal deletion mutant of HMGN1 (HMGN1-ΔC) and polyacrylamide gels of purified HMGN1 and the deletion mutant; right, quantification of mobility shift of acetylated and non-acetylated rMN with the HMGN1-ΔC particles. (**C).** Linker DNA is necessary for the preferential binding of HMGN to acetylated nucleosomes. Left: Schematic diagram of rMN and nucleosome core particle (rCP); center: gels of DNA from acetylated and non-acetylated rMN and rCP particles; left bottom: images of two-color mobility shifts. Right, quantification of two-color gel mobility shift assays of HMGN1 with H3K27ac and nonacetylated rMN and rCP. Error bars show standard deviation, n=2.

**Supplemental Figure S5.**
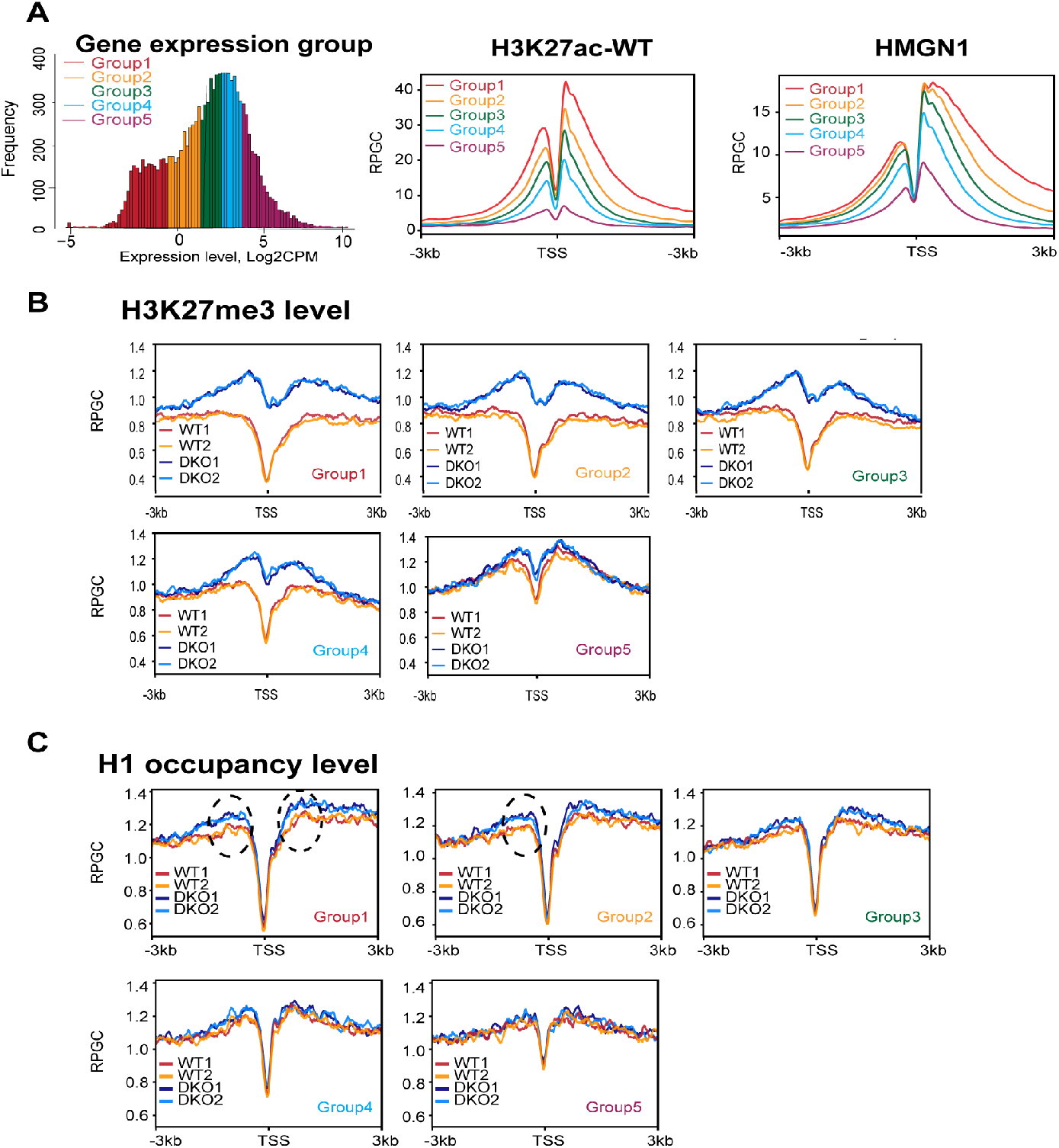
HMGN mediated epigenetic changes correlate with the magnitude of gene expression. **(A)** Genes were sorted into 5 tiers according to expression levels (left). Both the H3K27ac levels (center) and the HMGN occupancy (right) at TSS correlate positively with gene expression levels. **(B)** H3K27me3 levels at the TSS of WT and DKO MEFs sorted by transcription level tier (as defined in panel A). Note that the largest difference between WT and DKO is seen at the most highly expressed genes, which have the highest content of H3K27ac and HMGN. **(C)** H1 occupancy levels at the TSS of WT and DKO MEFs sorted by transcription level tier. Note that the largest difference between WT and DKO is seen at the most highly expressed genes. All ChIP analyses done with 2 biological replicates.

**Supplemental Figure S6.**
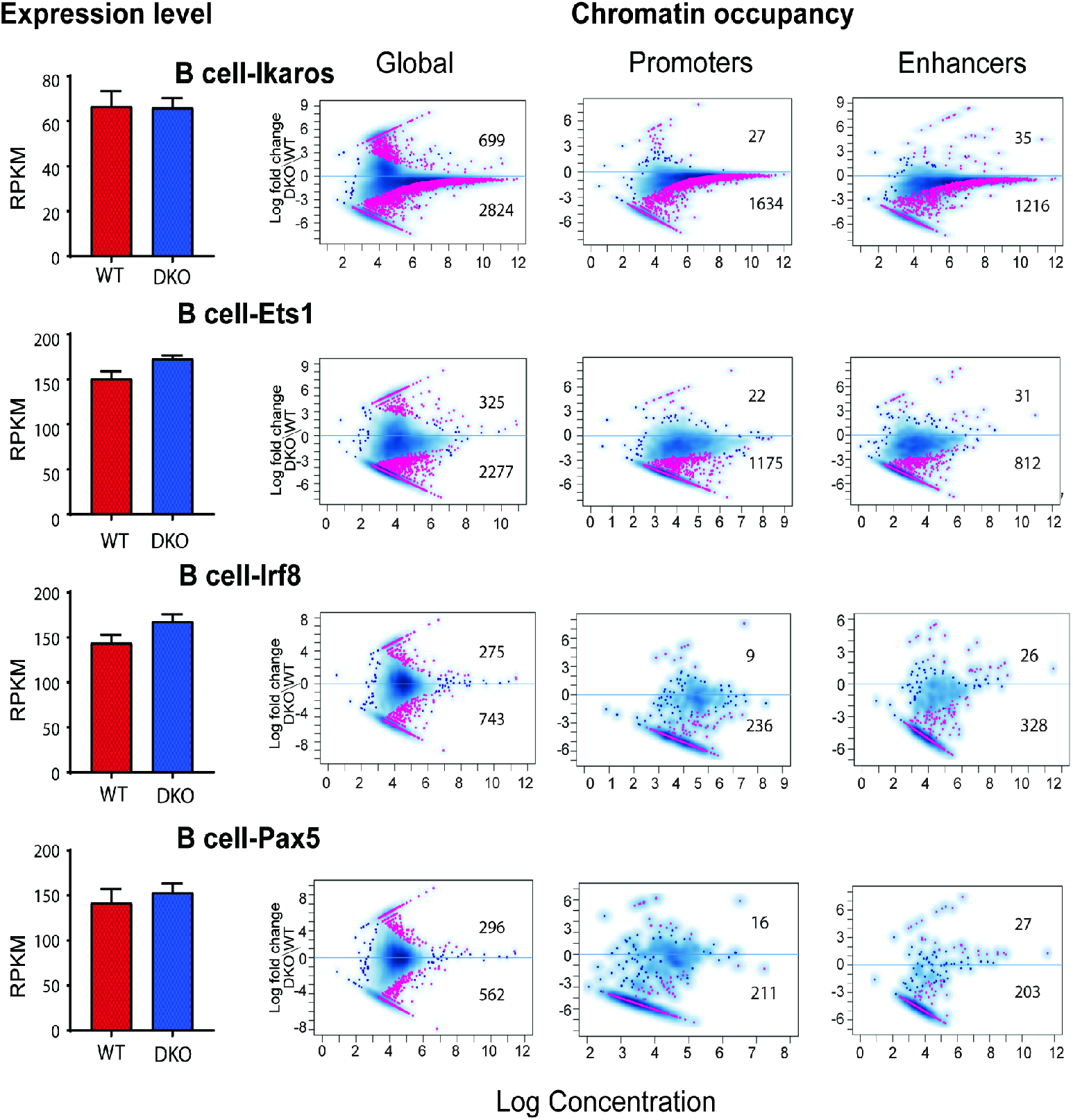
Altered chromatin occupancy of transcription factors in DKO resting B cells. Bar graphs show transcript levels determined by RNA seq analysis. MA plots show differences in TF chromatin binding between DKO and WT cells. Statistically significant differences (FDR<0.05) are shown in red. Blue dots and blue density cloud represents all points corresponding to the non-changing regions. All data from 2 biological replicates.

## Notes

### Competing Interest Statement

The authors have declared no competing interest.

